# Culturomics-Guided Precision FMT Enables Stable Antibiotic-Independent Engraftment via Metabolic Niche Targeting

**DOI:** 10.64898/2026.10.25.761671

**Authors:** Daniel M. Ferrer, Mark Holton, Minshen Yang, Ileana Villamil, Guillaume Urtecho

**Affiliations:** Department of Medicine, Division of Gastroenterology, University of California, San Diego, La Jolla, CA, USA; Biomedical Sciences Graduate Program, University of California, San Diego, La Jolla, CA, USA

**Author notes:** Corresponding author: Guillaume Urtecho, PhD, 9500 Gilman Dr, MC 0983, University of California, San Diego, La Jolla, CA 92093-0983, USA. These authors contributed equally to the work (Daniel M. Ferrer, Mark Holton).

**Keywords:** fecal microbiota transplantation, culturomics, metabolic niche profiling, colonization resistance, antibiotic-independent engraftment, donor-recipient complementarity, niche-associated communities, gut microbiome

## Abstract

The gut microbiome’s functional architecture can be conceptualized as a network of metabolic niches, yet clinical approaches to fecal microbiota transplantation (FMT) do not leverage niche - level information to guide engraftment. Here, we present a culturomics-based framework that profiles the metabolic niche landscape of individual microbiomes to enable targeted, antibiotic-independent FMT. Using a panel of 60 diverse media conditions, we show that gut communities from genetically identical mice harboring distinct microbiomes exhibit reproducible, vendor - specific niche-occupancy profiles, with each condition selecting for compositionally distinct niche-associated communities (NACs). We demonstrate that antibiotic pretreatment prior to FMT compromises community resilience, yielding communities that lose donor taxa more rapidly and recover less taxonomic diversity after secondary perturbation compared to untreated recipients. By comparing donor and recipient niche profiles, we identify vacant metabolic niches in recipient microbiomes and show that NACs targeting these vacancies achieve stable engraftment persisting for over 72 days without antibiotic pretreatment, whereas NACs targeting occupied niches fail to colonize. Furthermore, complex NACs not only fill vacant niches but reshape the surrounding native community, while simpler NACs engraft with minimal collateral impact. This culturomics-guided framework provides a scalable, mechanistic approach for rational donor-recipient matching that could advance personalized microbiome-based therapies.

## INTRODUCTION

The gut microbiome is a dense and dynamic ecosystem whose compositional stability and functional capacity profoundly shape host physiology, impacting nutrition, immunity, and disease risk ^1–3^. Under homeostatic conditions, thousands of bacterial taxa coexist in the gastrointestinal tract, harvesting otherwise inaccessible nutrients, synthesizing essential metabolites, and conferring protection against pathogens ^1,4^. Yet, subtle perturbations such as antibiotic exposure, dietary shifts, or xenobiotic insults can destabilize this balance, leading to dysbiosis and contributing to conditions such as inflammatory bowel disease, metabolic syndrome, and obesity^5^; ^6,7^. Therefore, there is a pressing need to understand and manipulate the microbiome as an essential component of our health.

The current gold standard for modifying the gut microbiome is fecal microbiota transplantation (FMT), in which a consortium of microbes from a healthy donor is administered to a patient with the goal of displacing or correcting a dysbiotic community. FMT has achieved remarkable success in treating recurrent *Clostridioides difficile* infections, but outcomes for other conditions such as inflammatory bowel disease and metabolic syndrome are inconsistent ^8,9^. Several factors limit the reliability of FMT as a general therapeutic. Engraftment outcomes are difficult to predict from patient to patient ^10–13^, and clinical practices vary widely across centers in route of administration, preparation of the fecal material, dosing schedule, and other parameters^12–14^. In many protocols, antibiotics are administered prior to transplantation to clear the endogenous microbiome and reduce competition with incoming donor taxa ^14–17^. However, antibiotic pretreatment itself influences the gut environment in ways that inhibit the formation of stable microbial communities, as the reduced microbial biomass promotes an aerobic gut environment that may eliminate strict anaerobes and thereby disrupt cross-feeding networks ^17^. Collectively, these variables contribute to the unpredictable efficacy of FMT and underscore the need for more ecologically centered strategies to manipulate the gut microbiome.

In order for an introduced microbe to stably integrate within a community, it must establish itself in an ecological niche. The gut ecosystem comprises many diverse metabolic niches, each defined by the nutrients and environmental conditions in a given region of the gut that support a subset of microbes capable of utilizing those resources^18^. However, niche availability is governed not only by resource landscapes but also by competition with the native microbiota ^19^. Competitive exclusion, in which a resident species prevents the engraftment of an incoming competitor, is a major barrier to microbial integration and has been most commonly observed between taxonomically related species that share overlapping resource requirements, rather than direct antagonism^20^. Consistent with this, donor-recipient complementarity, where the donor harbors species that are evolutionarily and metabolically different from the recipient’s residents, is among the strongest predictors of FMT success ^21^. The mechanistic basis of complementarity is likely multifactorial, but the ability to predict FMT outcomes from donor-recipient dissimilarity implicates niche-level competition as a major determinant^21,22^.

If the metabolic niches of a recipient’s microbiome could be profiled prior to transplantation, donors or defined communities could be selected to colonize unoccupied niches rather than compete for already-occupied ones. The power of targeting vacant niches is evident in studies showing that by creating an exclusive niche, exogenous microbes may be introduced and persist long term ^23,24^. However, it stands to reason that within any given microbiome there may exist niches that are underutilized or transiently available, yet no current approaches can systematically identify such vacancies or predict when they arise. Current experimental methods typically screen microbes against panels of gut-associated nutrients or defined metabolic traits to characterize specific niches, such as polysaccharide metabolism, but these assays do not reflect the complex molecular profiles of niches in vivo^25,26^. Computational approaches infer niche structure from genetic overlap or predicted metabolic capabilities of community members ^27,28^, yet may miss interactions that are not apparent in silico. Notably, both experimental and computational strategies tend to characterize niches at the level of individual isolates rather than complex communities, which can access niches through secondary or cooperative interactions. A more comprehensive framework for profiling how diverse communities occupy molecularly distinct niches within individual gut environments is therefore needed to enable precision microbiome therapeutics.

In this work, we developed a culturomics-guided framework for profiling occupied metabolic niches within microbiomes that enables targeted, antibiotic-independent transplantation into vacant niches. We show that antibiotic pretreatment prior to FMT produces communities that are less resilient to secondary perturbation, exhibiting greater compositional disruption and slower recovery compared to communities that received FMT without pretreatment. To perform antibiotic-independent transplantation, we leverage a panel of 60 diverse media conditions, each representing distinct *in vivo* niches, to profile which niches are functionally accessible by an individual’s microbiome. We then show that vacancies in these niche profiles can be targeted for transplantation by complementary niche-associated communities (NACs) and stably engraft for over 72 days. Moreover, we show that NACs engrafted in this manner can induce predictable shifts in the metabolome of these mice, inferred by metagenomically profiling these communities prior to transplantation. Together, this work lays the foundation for more precise FMT therapies that provide unprecedented customization of an individual’s gut microbiome to enhance their health.

## RESULTS

### Antibiotic pretreatment reduces the microbiome’s stability and resilience to future **perturbations**

While antibiotic pretreatment is an effective way to reduce the effects of colonization resistance to transplanted communities, the resulting increase in luminal oxygen has been shown to inhibit the growth of strict anaerobic microbes. We hypothesized that antibiotic pretreatment disrupts the obligate anaerobic taxa that anchor community stability, rendering the community more susceptible to subsequent perturbations. To test this, we compared the composition and stability of gut communities in mice that received antibiotics prior to FMT versus those that did not (**Figure 1A**). In prior work, we showed that stool communities derived from Envigo (Harlan) C57BL/6 mice (hereafter Env) could robustly colonize other microbiomes in the absence of any antibiotic pretreatment^29^. We therefore acquired C57BL/6 mice from Jackson Laboratories (hereafter Jax) and split them into two groups (n=4 each): one received an oral antibiotic cocktail of ampicillin, metronidazole, neomycin, and vancomycin for eight days (Jax^Abx^) to deplete the native microbiome, while the other received regular water (Jax). Antibiotic depletion was confirmed by qPCR, with a ΔCq of 8.05 between pre-and post-antibiotic samples corresponding to a 175–265-fold reduction in bacterial biomass (**Supplementary Figure 1**). Both groups then received a single dose of FMT from Env mice. 20 days after FMT, we assessed community resilience by administering the same antibiotic cocktail to Jax, Jax^Abx^, and Env donors for four days and then longitudinally monitoring the microbiome over the subsequent 11 days.

**Figure 1.**
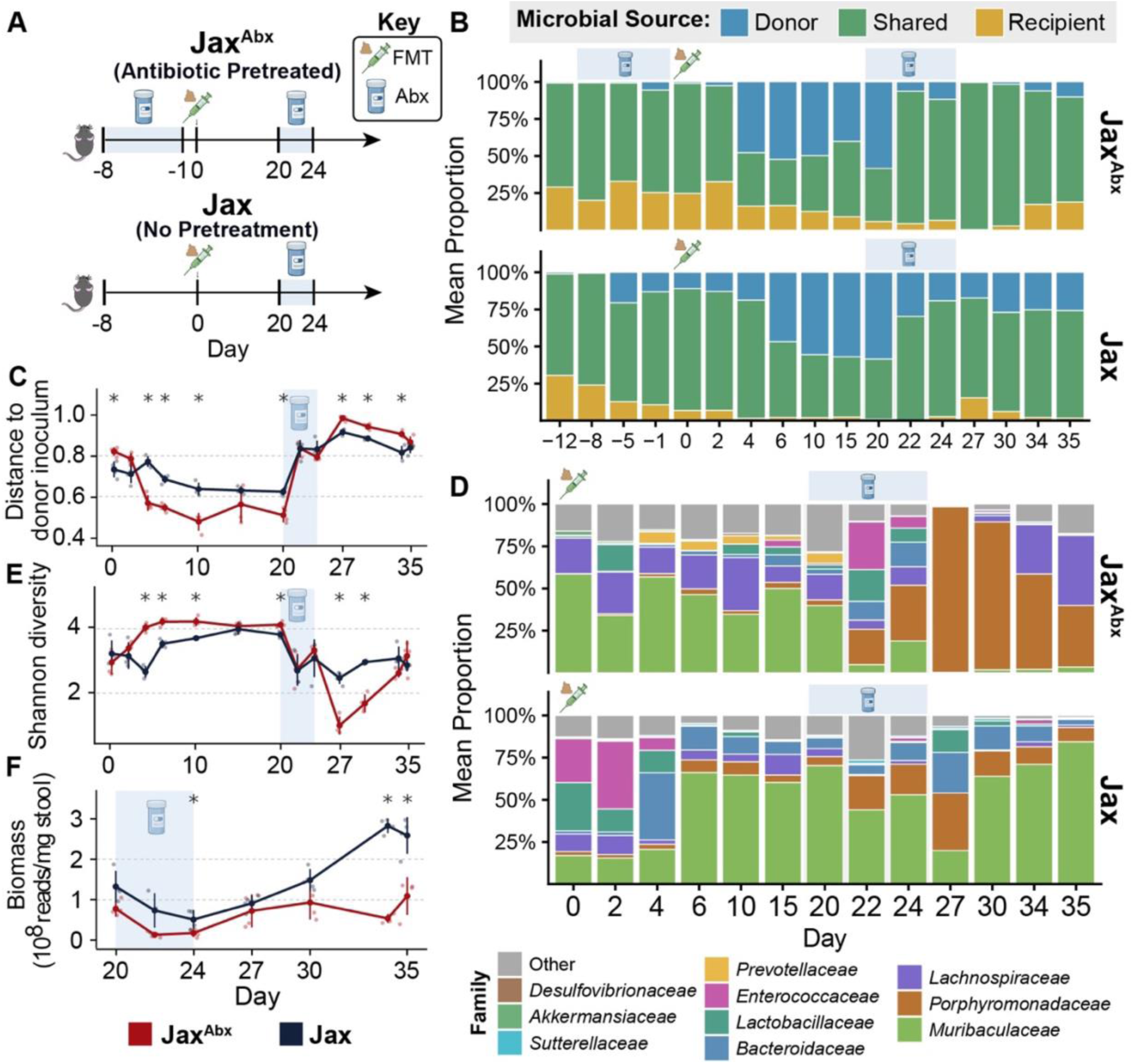
Antibiotic pretreatment reduces the microbiome’s stability and resilience to future perturbations. **A)** Experimental design. C57BL/6 mice from Jackson Laboratories (Jax) were split into two groups (n=4 each). One group received an oral antibiotic cocktail (ampicillin, metronidazole, neomycin, and vancomycin) for seven days (Jax^Abx^), while the other received regular water (Jax). Both groups received a single fecal microbiota transplant (FMT) from Envigo (Env) donor mice. Twenty days post-FMT, all groups including Env donors received a four-day secondary antibiotic challenge, followed by 11 days of recovery monitoring. **B)** Relative abundance of OTUs classified as donor-exclusive (blue), shared (green), or recipient-exclusive (yellow) across the time course for Jax^Abx^ (top), Jax (middle), and Env donors (bottom). Each bar represents group average at a given time point. **C)** Community similarity to the donor microbiome over time for Jax^Abx^ and Jax groups. Following FMT, Jax^Abx^ communities were initially more similar to the donor, but this pattern reversed after the secondary challenge, with Jax communities maintaining greater donor similarity through recovery. **D)** Family-level taxonomic composition across the time course for Jax^Abx^ (top) and Jax (bottom). **E)** Shannon diversity index over time for Jax^Abx^ and Jax groups. **F)** Total bacterial biomass during the recovery period following the secondary antibiotic challenge. * = p < 0.05 (Wilcoxon Rank Sum Test).

To track the fate of engrafted taxa, we performed longitudinal 16S V4 sequencing of stool samples from all mouse groups and monitored the dynamics of OTUs that were exclusively found in the donor, native to the recipient, or shared by both (**Figure 1B**). The Jax and Jax^Abx^ groups achieved similar distributions of donor, recipient, and shared microbes following FMT, although the transfer occurred over a slower time scale for Jax compared to Jax^Abx^ mice. Following the secondary antibiotic challenge, however, the two groups diverged sharply. Jax^Abx^ mice exhibited near-complete loss of donor microbes by day 27, retaining only 0.4% donor-derived OTUs, whereas Jax mice retained a donor fraction of 25.8% at day 35 (**Figure 1B**). We additionally assessed community similarity to the donor microbiome across the time course (**Figure 1C**). Following FMT, Jax^Abx^ communities were initially more similar to the donor, consistent with the more rapid and complete displacement of recipient taxa by antibiotic clearance. However, following the secondary antibiotic challenge, this pattern reversed, with Jax^Abx^ communities diverging from the donor profile while Jax communities maintained greater donor similarity through the recovery period.

We next examined the taxonomic composition of these populations throughout the time series (**Figure 1D**). Post-FMT, Jax^Abx^ communities were dominated by *Muribaculaceae* (42.7%) and *Lachnospiraceae* (19.2%), while Jax communities originally contained *Enterococcaceae* (26.0%) and *Lactobacillaceae* (28.6%) that rapidly declined as *Bacteroidaceae* and *Muribaculaceae* expanded. The secondary antibiotic challenge triggered distinct recovery trajectories in the two groups. By day 27, *Porphyromonadaceae* accounted for 98.1% of reads in Jax^Abx^ mice, whereas Jax mice still exhibited more diverse populations (**Figure 1D**) (Mann-Whitney U Test, *p* = 0.029). Shannon diversity followed a parallel pattern, with Jax^Abx^ communities exhibiting higher diversity immediately after FMT but lower diversity than Jax communities following the perturbation (**Figure 1E**). Jax^Abx^ mice also showed impaired biomass recovery following the challenge, with Jax communities reaching 5.3-fold higher total biomass by day 34 (**Figure 1F**). Taken together, these data indicate that antibiotic pretreatment prior to FMT yields communities that are less resilient, and subsequent perturbation substantially disrupts community composition, diversity, and donor engraftment.

### Culturomics reveals niche occupancy and niche-associated communities in the gut

Having established that antibiotic pretreatment compromises community resilience and transplant stability, we sought to develop antibiotic-independent approaches to allow for the stable transfer of gut microbiota. Rather than depleting resident communities to create open niches for donor engraftment, we considered whether donor communities could instead be directed into niches already unoccupied by the recipient microbiome. To test this, we asked whether metabolic niches could be profiled in vitro to guide targeted engraftment without antibiotic clearance. We reasoned that culturing stool communities in media replicating diverse nutrient conditions would simulate certain metabolic niches in the gut; and that conditions supporting growth indicate occupied niches, while those failing to support growth suggest vacant or underrepresented niches. Comparing donor and recipient growth patterns across such a panel would reveal which recipient niches are unoccupied and which donor-derived niche-associated communities (NACs) could fill them.

We assembled a curated panel of 60 media conditions drawing on the work of the Mouse Intestinal Bacterial Collection^34^, which successfully cultured microbial isolates representing 60-90% of the functional diversity detected in gut metagenomes. This panel consisted of a broad collection of media ranging from generic broths (e.g., Anaerobe Basal broth, Brain Heart Infusion broth) to specialized anaerobic formulations (e.g., Schaedler broth, Wilkins-Chalgren Anaerobe broth) and substrates that enrich for particular functional groups (e.g., *Akkermansia*-enriched media, Yeast Casitone Fatty Acids broth). By leveraging this diverse media collection, we aimed to extract specific subsets of the microbiome corresponding to distinct metabolic niches, thereby revealing which niches are occupied or vacant in a given community.

Initial profiles using a representative murine microbiome showed substantial variation in growth across the 60 media conditions, with some substrates supporting robust community expansion and others yielding minimal biomass (**Supplementary Figure 2**). To assess the reliability of this approach, we inoculated biological replicates from independent stool samples and observed high reproducibility between repeat experiments, indicating that niche-associated growth profiles are stable properties of a given community (**Supplementary Figure 3**). We next compared the total number of taxa captured across the full media panel to those detectable in stool by standard 16S V4 sequencing. Collectively, the 60 culture conditions selected for distinct taxonomic profiles across media (**Supplementary Figure 4**) and recovered more taxa at greater than 0.1% relative abundance than were observed in stool alone (**Supplementary Figure 5**), suggesting that in vitro niche profiling can reveal community members that fall below detection limits in bulk sequencing but become apparent when enriched under selective growth conditions.

We next asked whether this panel could capture niche-level variation between natural gut microbiomes. We acquired C57BL/6 mice (n=4 per vendor) from four vendors: Jackson Laboratories, Envigo, Charles River, and Taconic, which despite being genetically related, carry diverse microbiomes that vary in complexity and composition^29^. To profile the niches occupied by each microbiome, we separately inoculated fresh stool from each mouse vendor into the full media panel and incubated for 48 hours under anaerobic conditions (**Figure 2A**). Across the microbiomes, Charles River microbiomes exhibited the most growth, in 33 of 60 conditions (**Figure 2B**). This growth was most frequently observed in complex, nutrient-rich media and was most sparse amongst the defined minimal media supplemented with individual, diverse carbohydrate sources, suggesting that niche occupancy varies across nutrient complexity and that defined carbon sources may delineate metabolic niches that are differentially accessible across microbiomes.

**Figure 2.**
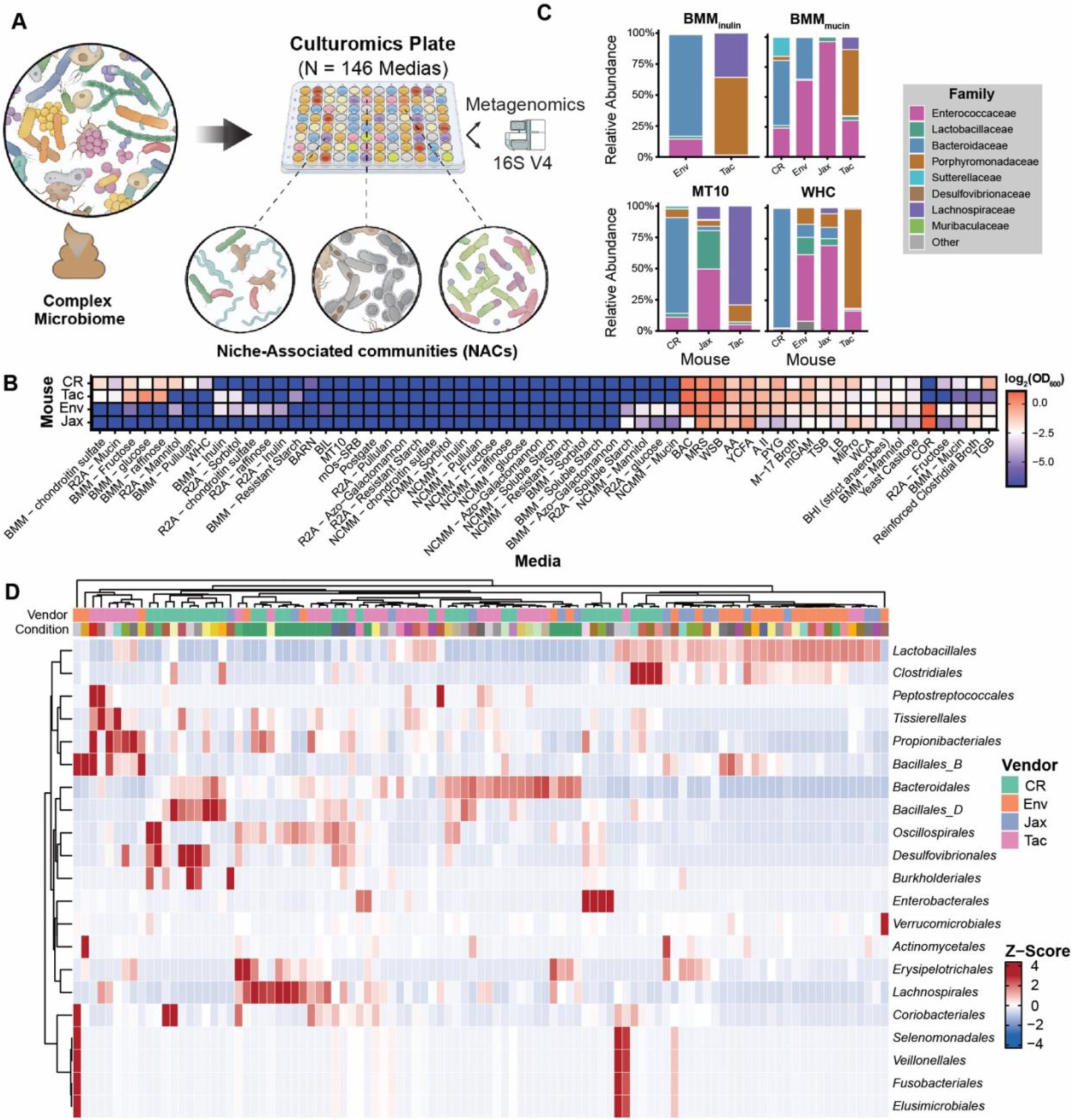
Culturomics reveals niche occupancy and niche-associated communities across diverse gut microbiomes. **A)** Schematic of the culturomics workflow. Stool from C57BL/6 mice sourced from four vendors (Jackson, Envigo, Charles River, and Taconic) was inoculated into a panel of 60 diverse media conditions and incubated for 48 hours under anaerobic conditions. Resulting niche-associated communities (NACs) were profiled by 16S V4 sequencing and metagenomics. **B)** Growth across the 60-medium panel for each vendor microbiome. Heatmap of log2-transformed growth values (Final OD_600_). **C)** Family-level taxonomic composition of NACs grown in selected media conditions (BMM_inulin_, BMM_mucin_, MT10, WHC) across vendors. **D)** Clustered heatmap of taxonomic order abundance (z-score) across all culture conditions and vendors. Rows represent bacterial orders and columns represent individual culture libraries, colored by vendor of origin. Hierarchical clustering reveals condition-specific and vendor-specific patterns of taxonomic enrichment across the panel.

To determine the composition of NACs growing under individual media conditions, we performed 16S sequencing across all vendor panels. NAC composition varied substantially both across media conditions and across source microbiomes cultured in the same condition (**Figure 2C**, **Supplementary Figure 6**). For example, BMM supplemented with inulin supported growth of Envigo and Taconic communities but not Jackson or Charles River, and the two communities that did grow differed markedly. Envigo communities were dominated by Bacteroidaceae (>75%) with minor populations of Enterococcaceae and Lactobacillaceae, whereas Taconic communities supported only Porphyromonadaceae and Lachnospiraceae. Similarly, MT10 media supported growth from Charles River, Jackson, and Taconic but not Envigo, and each was composed of distinct, complex populations of four to five detectable families per community. In contrast, BMM supplemented with mucin and WHC supported growth from all four vendor microbiomes, each yielding communities of similar family-level taxonomic composition, but different distributions of each. These results demonstrate that individual media conditions capture compositionally distinct NACs between microbiome inputs.

We next explored across all niches to evaluate the diversity and comprehensiveness of communities we could capture using this method (**Figure 2D**). Family-level analysis across conditions in different mice revealed the breadth of the microbiome captured by the culturomics approach. These data reveal the taxa composing communities grown in different media and the prevalence of these taxa across conditions. These data also reveal which taxa are frequently found in the same communities, a potential sign of beneficial interspecies relationships. Comparison of community diversity from different mice in the same media condition further reveals how NAC composition varies. Together, these results demonstrate that our culturomics panel captures diverse niche profiles of natural gut communities and reveals the taxonomic diversity within individual niches.

### Vacant niches can be targeted for precision engraftment by complementary donor communities

Having established that our culturomics panel captures niche-level variation between natural gut microbiomes, we next tested whether vacant niches in recipient microbiomes could be targeted for stable engraftment without antibiotic pretreatment. We hypothesized that individual NACs derived from a donor could be transplanted into a recipient, and those complementary to unoccupied niches would engraft and persist, whereas NACs targeting occupied niches would be competitively excluded. To test this, we profiled Envigo (Env) and Jackson Laboratories (Jax) mice using our culturomics panel, growing stool samples under anaerobic conditions, and identified conditions with differential growth between communities (**Figure 3A**). We selected two conditions in which donor mice displayed NAC growth but recipient mice did not, representing putatively vacant niches: BAC and BMM supplemented with mucin (BMM_muc_) (**Figure 3B**). We also included YCFA, a condition in which both donor and recipient communities grew, representing an already occupied niche in the recipient. 16S sequencing of the donor NACs prior to gavage revealed varying levels of richness (BAC: eight OTUs; BMM_muc_: four OTUs; YCFA: two OTUs). We grew each donor NAC, individually administered them to Jax recipients via oral gavage (n = 4 per NAC), and compared engraftment against PBS-gavaged controls.

**Figure 3.**
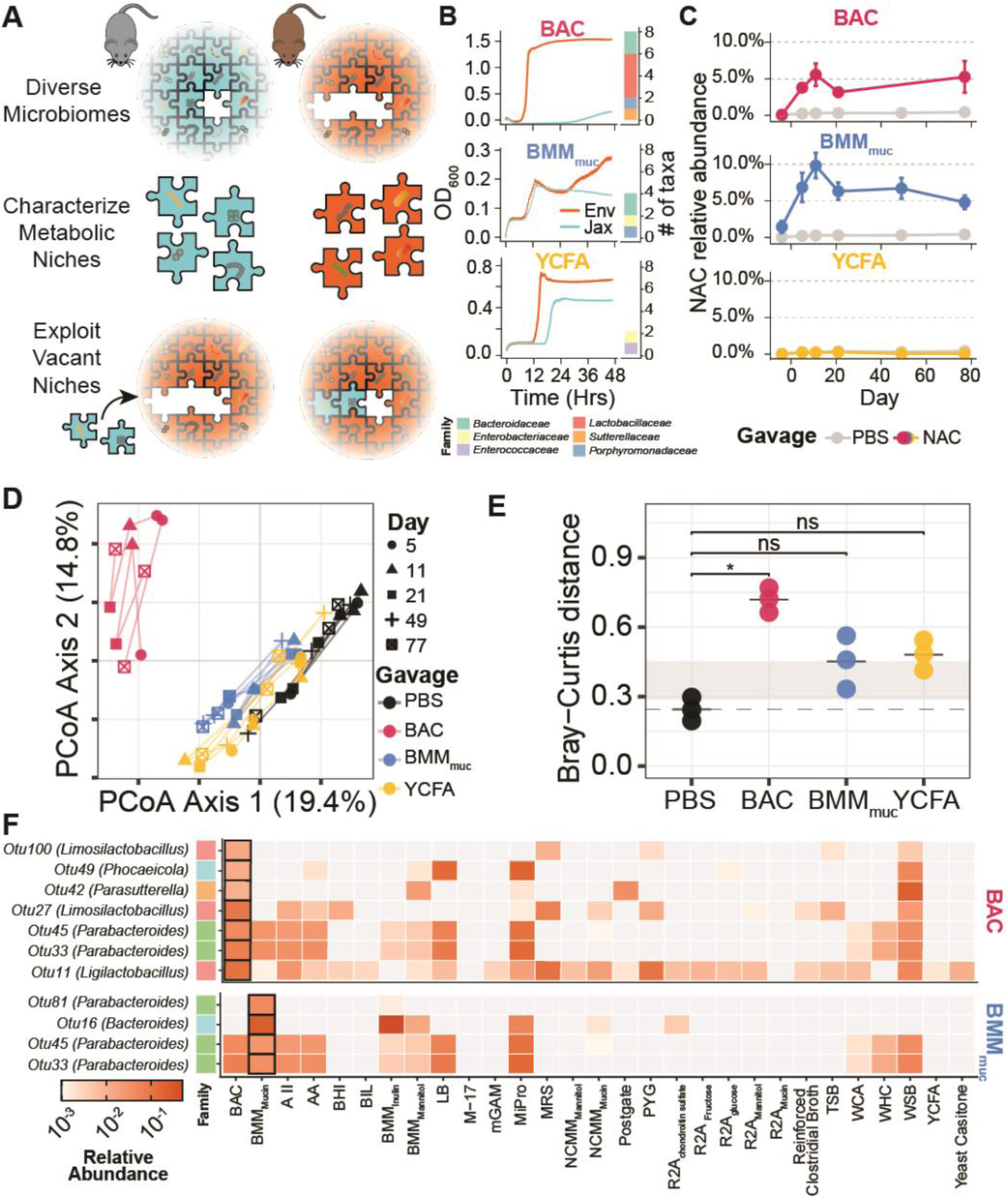
Vacant niches can be targeted for precision engraftment by complementary donor communities. **A)** Schematic of the niche-targeted transplantation strategy. Microbiomes from C57BL/6 mice with distinct community compositions are profiled using the culturomics panel to identify metabolic niches. Vacant niches in the recipient are identified and targeted for engraftment with donor-derived NACs. **B)** Growth curves (OD_600_ over 48 hours) and family-level composition of Envigo (Env, orange) and Jackson (Jax, light blue) stool cultured in BAC, BMM_muc_, and YCFA media. Marginal stacked bars show counts of unique OTUs identified in the resulting NACs, colored by family-level composition. **C)** Longitudinal relative abundance of NAC-associated taxa in recipient Jax mice over 73 days post-gavage. NAC-gavaged mice (colored) are compared to PBS controls (gray). BAC and BMM_muc_ NACs persisted throughout the experiment, while YCFA NAC taxa were not detected at any time point, consistent with competitive exclusion from an occupied niche. Error bars represent standard error of the mean. **D)** Principal coordinates analysis (PCoA) of Bray-Curtis distances for recipient communities over time, with NAC-derived OTUs computationally masked to isolate changes in the native community. Points are colored by gavage group (PBS, BAC, BMM_muc_, YCFA) and shaped by collection day. **E)** Bray-Curtis distance between post-gavage and pre-gavage community profiles for each treatment group, with NAC taxa masked. Only the BAC NAC produced a significant compositional shift relative to PBS controls (*p < 0.05, Wilcoxon rank-sum test). The dashed line and gray band represent the mean and range of PBS control distances. ns, not significant. **F)** Heatmap of relative abundance for BAC (top) and BMM_muc_ (bottom) NAC OTUs across 29 culture conditions in the culturomics panel. OTUs are ordered by decreasing individual niche breadth (number of conditions detected). Color intensity reflects log_10_-scaled relative abundance (orange gradient); gray cells indicate conditions in which the OTU was not detected. Black-bordered cells denote the source culture condition from which each NAC was derived. Colored bars at left indicate bacterial family assignment.

Five days post-gavage, we cultured recipient stool from each group in the media condition from which their respective NAC was derived. While PBS control stool still showed no growth in BAC or BMM_muc_, all three NAC-gavaged groups produced significant growth in their respective conditions, confirming successful engraftment of donor-derived taxa into vacant recipient niches (**Supplementary Figure 7**). To assess whether engraftment was stable, we performed longitudinal 16S sequencing over 73 days to track the abundance of microbes corresponding to each Envigo NAC (**Figure 3C**). NACs targeting the two vacant niches, BAC and BMM_muc_, persisted in recipient gut communities for the duration of the experiment, consistently comprising 3.7% and 12.1% of the total community, respectively. In contrast, microbes derived from the Envigo YCFA NAC were not detected in recipient communities at any time point during longitudinal monitoring, indicating that it failed to engraft. The increased growth observed for YCFA at five days post-gavage was likely attributable to a bloom in endogenous Jax taxa rather than to donor-derived microbes, as PBS control stool cultured in YCFA medium showed initial growth at day five that declined by day 73, suggesting transient fluctuations in endogenous populations rather than stable colonization.

To distinguish whether NAC engraftment reshaped the native recipient community or whether transplanted taxa persisted without influencing the resident microbiome, we profiled the recipient native communities longitudinally. To isolate changes in the native community from the direct contribution of transplanted taxa, we computationally masked all NAC-derived OTUs and reanalyzed community composition. PCoA of these masked profiles revealed that PBS controls and YCFA-gavaged mice overlapped substantially across time points, indicating that YCFA gavage did not alter the native community beyond baseline variation (**Figure 3D**). In contrast, BAC-gavaged mice exhibited the greatest separation from controls even after NAC taxa were removed, and BMM_muc_-gavaged mice showed intermediate separation (**Figure 3D**). Beta diversity analysis comparing masked post-gavage to pre-gavage baselines confirmed that only the BAC NAC produced a significant compositional shift relative to PBS controls (Wilcoxon rank-sum test), while BMM_muc_ did not reach significance (**Figure 3E**). These results suggest that NAC engraftment may reshape the native community with the magnitude of this effect varying by NAC composition.

We next asked what determines the magnitude of community reorganization among NACs that successfully engraft. Both BAC and BMM_muc_ NACs achieved stable colonization, yet BAC drove significantly greater compositional changes in the native community despite comprising a smaller fraction of the total community than BMM_muc_. We hypothesized that this disparity reflects differences in the metabolic versatility of their constituent taxa and that BAC taxa may expand into other niches in the gut after establishing themselves. To test this, we analyzed the niche breadth of BAC and BMM_muc_ OTUs across our culturomics panel (**Figure 3F**). Although individual BAC and BMM_muc_ OTUs did not differ significantly in the number of conditions in which they were detected (BAC median: 7, range: 2–16; BMM_muc_ median: 6, range: 1–8; Mann-Whitney U, p = 0.77), the BAC NAC collectively spanned a far broader range of the culturomics panel. Together, the seven BAC OTUs were detected across 22 culture conditions, whereas the four BMM_muc_ OTUs spanned only 10 conditions (**Figure 3F**). This difference reflected both the larger number of constituent taxa in the BAC NAC and their phylogenetic and metabolic diversity, with each OTU occupying distinct profiles niches. The broad collective niche breadth of BAC taxa is consistent with the capacity of these organisms to expand beyond the BAC niche into additional metabolic niches, thereby influencing community composition across diverse microenvironments. By contrast, BMM_muc_ taxa occupied a narrower, more specialized set of conditions, consistent with stable engraftment that produces limited community-wide effects.

Taken together, these results demonstrate that niche-targeted NAC transplantation achieves stable, long-term engraftment without antibiotic pretreatment, and that the degree of community-level reorganization scales with the collective metabolic breadth of the transplanted community rather than its relative abundance.

## DISCUSSION

In this study, we developed a culturomics-guided framework for profiling metabolic niches in the gut microbiome and used it to improve the precision of FMT. We first demonstrated that antibiotic pretreatment, a common clinical practice intended to enhance engraftment, paradoxically compromised community resilience. Pretreated mice lost donor-derived taxa more rapidly, recovered less taxonomic diversity, and failed to regain peak biomass after a secondary perturbation. We then used a panel of diverse culture conditions paired with sequencing to profile niche occupancy across microbiomes from genetically identical mice sourced from different vendors, revealing distinct community architectures defined by which niches were filled and by which taxa. Finally, we leveraged these profiles to perform targeted transplantations into specific niches, demonstrating that NACs directed at vacant niches engrafted stably and persisted long-term, whereas those targeting occupied niches did not. Together, these results establish metabolic niche profiling as a strategy for achieving stable, antibiotic-independent engraftment.

Our findings add to a growing body of evidence that antibiotic pretreatment should be reconsidered in FMT protocols. In our experiments, both antibiotic-treated groups exhibited lower engraftment than the FMT-alone group, and pretreated communities showed reduced resilience to subsequent perturbation. These observations are consistent with prior work demonstrating that antibiotic pretreatment can reduce donor strain engraftment relative to FMT alone ^30^ and that the ecological consequences of antibiotic exposure persist for months, with key taxa remaining depleted for up to eight months after treatment ^31,32^. The durability of antibiotic-induced dysbiosis underscores the need for engraftment strategies that do not rely on clearing the recipient community. Our results demonstrate that such strategies are feasible, as donor communities can establish in vacant metabolic niches without antibiotic-mediated niche clearance, preserving the stability of the existing community.

Our results suggest that metabolic niche availability is a principal determinant of engraftment success, a framework that unifies several observations from the FMT literature. The concept of “super donors,” whose stool achieves successful FMT more frequently, has been attributed to high community richness and favorable host factors ^33,34^, and studies have shown that richer donor communities can colonize less rich recipients but not the reverse^35^. We propose that these observations reflect niche-level dynamics. Communities with greater taxonomic richness and phylogenetic diversity likely encompass a broader range of metabolic capabilities, allowing them to fill more recipient niches upon transplantation. This interpretation aligns with recent ecological analyses of FMT outcomes. Schmidt *et al.* examined metagenomes from 316 FMTs and identified donor-recipient complementarity as a key predictor of engraftment, defining it as the degree to which donor strains are phylogenetically or metabolically distinct from the recipient community ^21^. Similarly, Pinto *et al.* found that ulcerative colitis patients who achieved sustained clinical response retained recipient strains while selectively incorporating complementary donor taxa, rather than experiencing wholesale community replacement ^36^. A complementary study using metabolic modeling found that optimal donors had minimal metabolic overlap with their recipients ^22^. Engraftment succeeds when donor organisms fill unoccupied niches rather than compete for resources already claimed by resident taxa.

Beyond explaining patterns of FMT engraftment, metabolic niche profiling opens a path toward precision microbiome therapies. Conventional FMT transfers the entire donor community without knowledge of which components will persist or what functions they will confer, limiting both the predictability and specificity of the intervention. By contrast, our platform enables identification of specific NACs that are compatible with a given recipient’s niche landscape, allowing for targeted modifications to the microbiome with more predictable outcomes. Previous work has shown that defined microbial introductions, from single species to small consortia, can influence host physiology through mechanisms including immune modulation^37^, enhanced colonization resistance^38^, and altered metabolite production^39^. Our culturomics-guided approach provides a systematic framework for determining which communities will be compatible with a particular recipient before transplantation. Realizing the full potential of this strategy will require characterizing NACs beyond their taxonomic composition. Metabolomic profiling of NAC cultures could reveal the functional outputs of each community, enabling selection of consortia based on desired metabolic activities. Co-culture of NACs with mammalian intestinal cells would further connect community composition to effects on host physiology, moving the field from empirical FMT toward rationally designed microbiome interventions tailored to individual patients.

Several limitations of the current study should be noted. Our culturomics panel profiles niche occupancy under *in vitro* culture conditions, and the extent to which these *in vitro* niches correspond to functional niches *in vivo* remains an open question. Host factors, spatial structure along the gastrointestinal tract, and dynamic nutrient availability may reshape competitive landscapes in ways not captured by our panel. Additionally, while we demonstrated that NACs can stably colonize vacant niches, we have not directly compared this targeted approach with conventional antibiotic-pretreated FMT in a controlled experiment. Such a comparison will be necessary to quantify the advantage of antibiotic-independent engraftment over current protocols. Our transplantations also used cultured communities rather than synthetic, defined consortia, and determining whether rationally designed communities can achieve comparable engraftment will be an important next step. Finally, translation to human microbiomes, which harbor greater interpersonal variation and are shaped by diet, medication, and immune status, will require further investigation. Despite these limitations, our findings establish metabolic niche profiling as a tractable framework for guiding microbiome interventions and provide a foundation for developing personalized, precision approaches to FMT.

## METHODS

### Antibiotic Treatment

To deplete the native gut microbiome prior to FMT, mice received a broad-spectrum antibiotic cocktail administered *ad libitum* in drinking water for 7 days. The cocktail consisted of ampicillin (0.5 g/L), metronidazole (0.5 g/L), neomycin (0.5 g/L), and vancomycin HCl (0.25 g/L) dissolved in 3% sucrose to improve palatability. Antibiotic water was prepared fresh and replaced every 2 days to maintain drug stability, and bottles were wrapped in aluminum foil to protect light-sensitive components from photodegradation. Cages were changed every 2 days to prevent coprophagy-mediated re-inoculation of depleted taxa. Mice were weighed on days 0, 4, and 7 of antibiotic treatment, and water intake was monitored at each bottle change to confirm adequate consumption. Following the 7-day antibiotic course, mice received regular (non-antibiotic) drinking water for 1 day before proceeding to FMT or subsequent experimental manipulations.

### Culturomics Panel Assembly and Inoculation

Media used in the culturomics panel were made according to their recipes (see: Culture media) and aliquoted into 50 mL tubes and frozen at-20℃ for long-term storage. 2 mL of each medium used were allotted into a 96-well deep well plate to serve as a master plate to ensure consistent media conditions for each experiment. Each well of a 96-well plate received 200 µL of their respective media from the master deep well plate. 200 µL of sterile water was used to fill the wells along the outer edge of the plate that received no media to reduce evaporation. Stool acquired from mice was used as the inoculum. In an anaerobic chamber, four stool samples from cage-mates (n = 4 mice per cage) sourced from the same vendor were individually resuspended in 1 mL of anaerobic PBS in a 1.5 mL tube and mechanically disrupted using an inoculating loop. The resuspended stool was allowed to sit for one minute to allow the solid particles to collect at the bottom of the tube. 50 µL of each resuspended sample were diluted into 8 mL of anaerobic PBS. Each media well received 10 µL of this pooled stool suspension. Plates were covered with a Breathe-Easy sealing membrane (Sigma-Aldrich, Z380059) and grown anaerobically in a Cerillo Alto Plate Reader (Cerillo, ALT-1600) for 48 hours with shaking at 300 RPM.

## Culture media

### Media recipes are found in **Table S1**. All quantities are per litre of medium

#### Metagenomic Library Preparation

Metagenomic sequencing libraries were prepared as previously described ^29^ using a scaled-down Tn5 tagmentation-based Nextera library preparation protocol with 2 ng gDNA as input ^40^. Libraries were submitted to the UC San Diego Institute for Genomic Medicine Genomics Center for sequencing on an Illumina NovaSeq X Plus platform (2 x 150 bp) following the manufacturer’s instructions.

#### Metagenomic analysis

Paired-end FASTQ files were processed using a custom Snakemake workflow for read quality control and host depletion prior to taxonomic profiling using Woltka. Reads were quality-filtered and adapter-trimmed with fastp (v1.01) ^41^ and aligned against a mouse genome reference (GRCm39) with Bowtie2 (v2.5.4) ^42^ in very-sensitive local mode. Only read pairs that did not align to the host reference were retained (i.e., unmapped concordant pairs) for downstream analysis. Host-depleted read pairs were further depleted by classification with Kraken2 (v2.17.1) ^43^ against a Kraken2 database (k2_standard_08_GB_20250714). To generate final host-depleted FASTQs for downstream profiling, reads assigned to *Eukaryota* (NCBI taxid 2759) were excluded (including all descendant taxa) using extract_kraken_reads.py from the KrakenTools suite ^43^. Downstream community profiling was performed using the Qiita metagenomics platform ^44^ with a standard Woltka metagenomics workflow (Woltka v0.1.7) ^45^ using the Web of Life reference (WoLr2) ^46^ for taxonomic and functional assignment. Genome counts were analyzed in R (v4.5.1) as follows: Genome count tables from Qiita were filtered to OTUs at > 0.1% abundance in at least 10% of all samples (across all diets and treatments) leaving 157 genomes.

#### Fecal Microbiota gDNA extraction and 16S rRNA amplicon sequencing

Fecal samples were disrupted in PBS using mechanical shearing and microbial gDNA was extracted using a previously published protocol^40^. 16S V4 regions were amplified in a single step PCR using the modified 515f ^47^ and 806r ^48^ primers that appended dual indices and flow cell adapters for Illumina sequencing. Samples were amplified until approximately mid-exponential phase and then pooled together based on the fluorescence increase at the last cycle. Pooled samples were concentrated using a DNA Clean & Concentrator-5 (ZYMO D4003) and run on a 2% E-gel EX Agarose Gel (Thermo Fisher G401002). DNA bands at ∼390bp were excised from gel and purified using a Zymoclean Gel DNA Recovery Kit (ZYMO D4008) following the manufacturer’s instructions. Purified libraries were sequenced on Illumina MiSeq i100 platform (2×150 cycles).

#### 16S rRNA amplicon analysis and OTU clustering

16S V4 sequencing reads were analyzed using USEARCH v11.0.667 ^49^ as follows. First, paired-end reads were merged using “-fastq_mergepairs” using default settings. Merged reads were then filtered using “-fastq_filter” mode with arguments “-fastq_maxee 1.0-fastq_minlen 240” to retain merged reads with less than 1 expected base miscall and greater than 240bp. Reads passing quality control were deduplicated (-fastx_uniques), clustered into OTUs (-unoise3) at 100% identity, and quantified within clusters (-otutab) to generate an OTU count table. Taxonomy of OTUs was assigned using the Ribosomal Database Project classifier ^50^ trained with 16S rRNA training set 18 ^51^. OTU count tables were normalized to relative abundance and filtered for OTUs with greater than 0.1% relative abundance on average across all samples for downstream analysis. Bray-Curtis dissimilarities, PERMANOVA (adonis2), and diversity metrics were computed on these filtered relative abundances using the vegan package, and figures were generated with ggplot2 and cowplot. Analysis was performed in R using custom code available as a GitHub repository, https://github.com/urtecholabucsd/Ferrer_Holton_CulturomicsFMT.

#### Single Community Gavage Selection and Transplantation

Candidates for the single community gavage were determined by growing stool samples from Envigo and Jackson Labs mice in our culturomics panel for 48 hours. We looked for specific growth patterns: 1) High growth in the donor mice, no growth in the recipient mice; 2) Low growth in donor mice, no growth in recipient mice; 3) Growth in both the donor and recipient mice. We defined high growth as an OD_600_ greater than 1.0 and low growth as greater than 0.1, but less than 0.5. After selecting the communities for gavage, we grew samples of the donor stool in the chosen media overnight. 1 mL of each culture was washed thrice in anaerobic PBS and resuspended in 1 mL anaerobic PBS before passing through a 100 µm filter. 200 µL of this NAC suspension was gavaged into each member of a cage (n=4 mice per cage), with each cage receiving a separate NAC. Stool samples were collected from the recipient mice periodically during the experiment for longitudinal analysis.

## Availability of Data and Materials

The datasets generated and/or analyzed during the current study are available at https://github.com/urtecholabucsd/FMT.

## Use of Generative AI

The generative AI tool Claude (Opus 4.6/5/5.5 Anthropic) was used during the preparation of this manuscript for coding assistance, editorial proofreading, identification of internal inconsistencies between text and figures, and copy editing of the final draft. All scientific content, analysis, interpretation, and writing were performed by the authors. The authors reviewed and take full responsibility for the final manuscript.

## ACKNOWLEDGEMENTS

G.U. acknowledges support from the Howard Hughes Medical Institute (GT17584), the NIDDK-funded San Diego Digestive Diseases Research Center (P30 DK120515), the NIH Common Fund’s Faculty Institutional Recruitment for Sustainable Transformation (FIRST) program (NCI U54CA272220), and the NCCIH (R21AT013617).

---

**Supplementary Figure 1.**
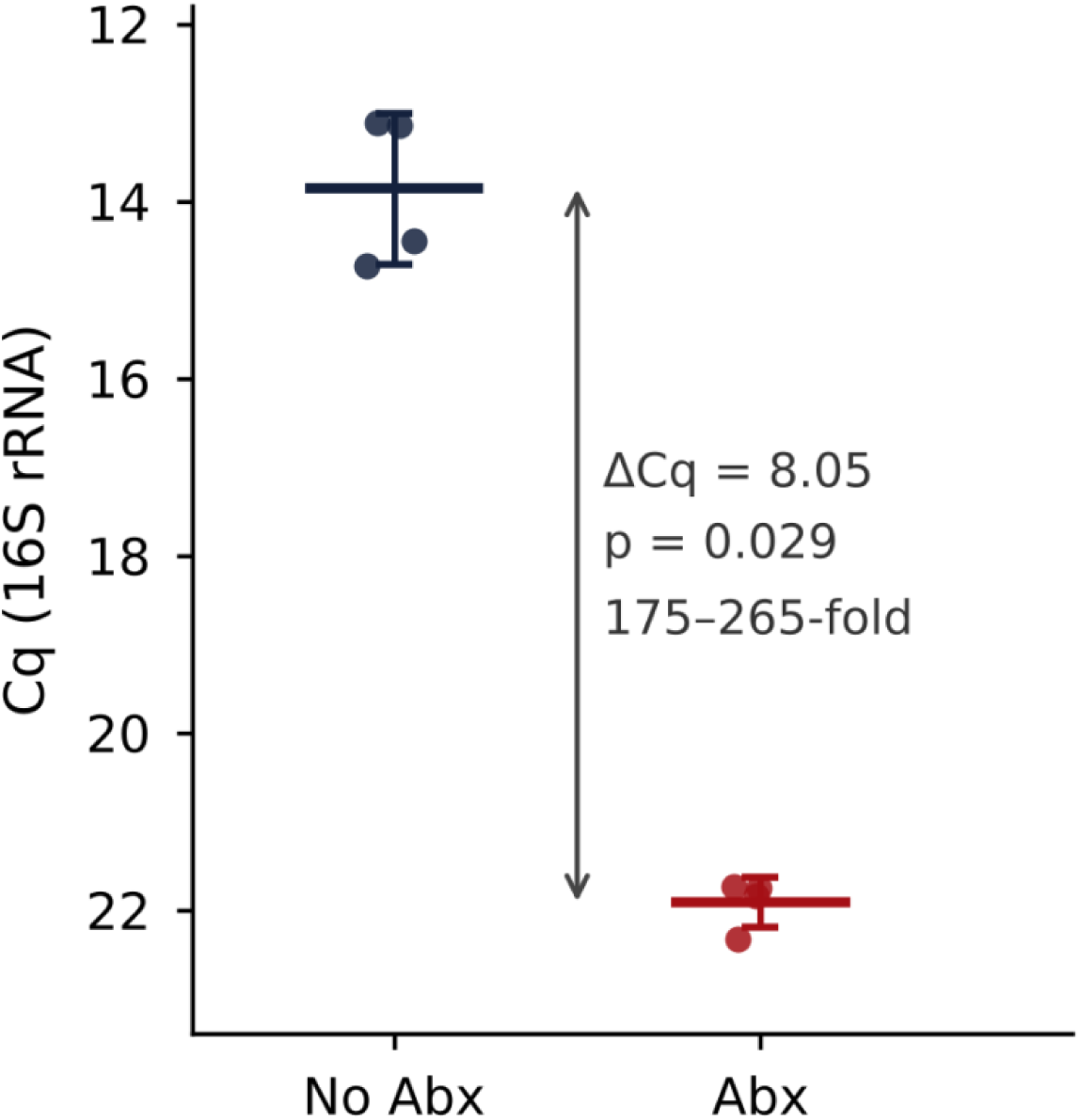
Antibiotic pretreatment depletes total bacterial load by over two orders of magnitude. Quantitative PCR of the 16S rRNA gene in stool from mice receiving the ampicillin, metronidazole, neomycin and vancomycin cocktail (Abx, n = 4) or regular water (No Abx, n = 4) for seven days. Points are individual mice, horizontal bars are group means and error bars are one standard deviation. Abx mice showed a mean Cq of 21.91 ± 0.28 compared with 13.86 ± 0.86 in untreated controls (ΔCq = 8.05; Mann-Whitney U, p = 0.029; Welch’s t-test, p = 1.1 × 10⁻⁴), corresponding to a 175-to 265-fold reduction in bacterial load for amplification efficiencies between 90% and 100%. A spike-in standard was run in parallel as a tenfold dilution series to quantify fold change. Y-axis is inverted to represent the inverse relationship with Cq and biomass.

**Supplementary Figure 2.**
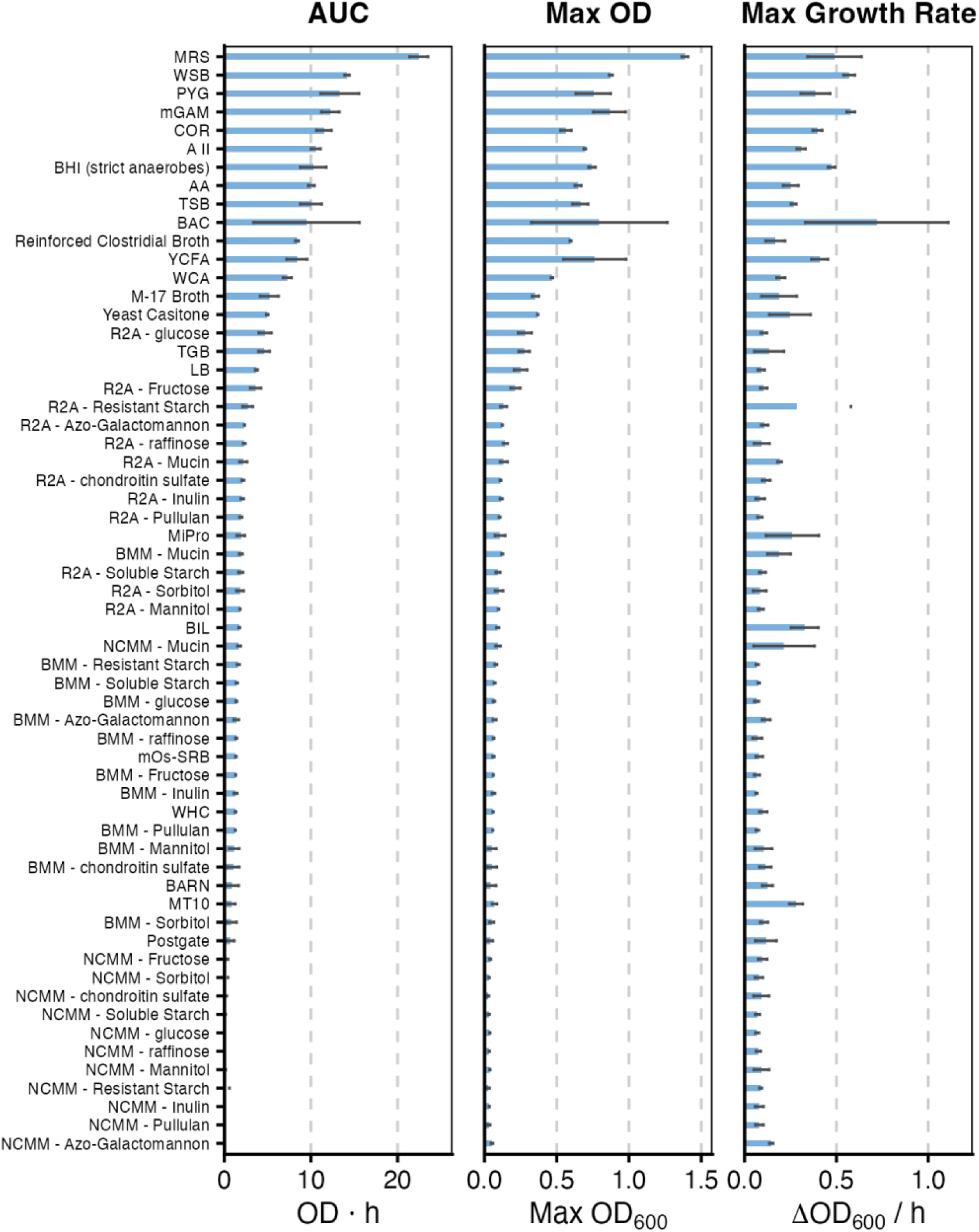
Growth metrics vary widely across the 60-medium panel. AUC (OD·h), maximum OD_600_, and μ (ΔOD_600_/h) are plotted for each medium, ranked by mean AUC across two Envigo replicates. Rich general-purpose media (MRS, mGAM, PYG, WSB) achieve the highest biomass, while minimal and substrate-supplemented media (BMM variants, NCMM, R2A) support substantially lower growth. This gradient reflects the selective pressure each medium imposes on the input community, with rich media promoting broad growth and defined media enriching for metabolically specialized subsets. Error bars represent variability between vendor replicates.

**Supplementary Figure 3.**
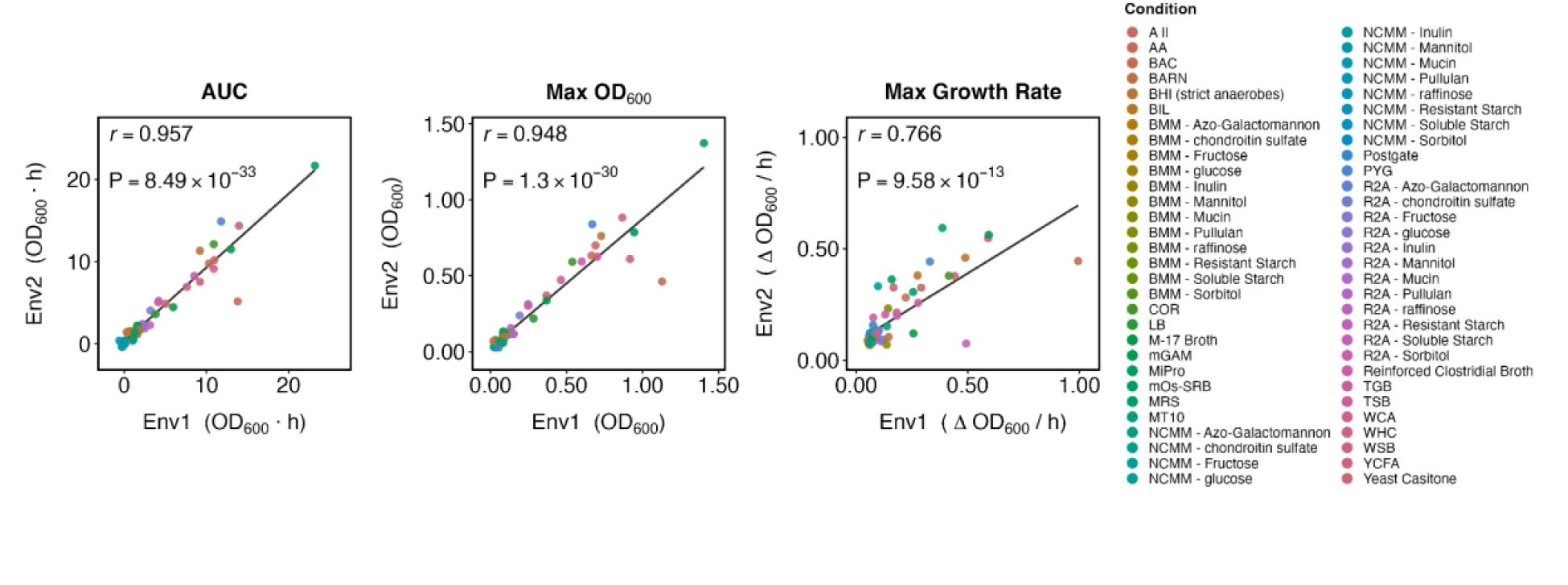
Growth metrics are reproducible across biological replicates. Scatter plots comparing AUC, maximum OD_600_, and μ between two independent Envigo stool inocula (Env1, Env2) cultured across all 60 media. AUC (r = 0.957) and maximum OD_600_ (r = 0.948) are highly correlated, while ΔOD_600_/h shows moderate correlation (r = 0.766). The high reproducibility of biomass metrics confirms that the panel produces deterministic growth responses from independent biological replicates. Pearson correlation coefficients are shown; each point represents one medium.

**Supplementary Figure 4.**
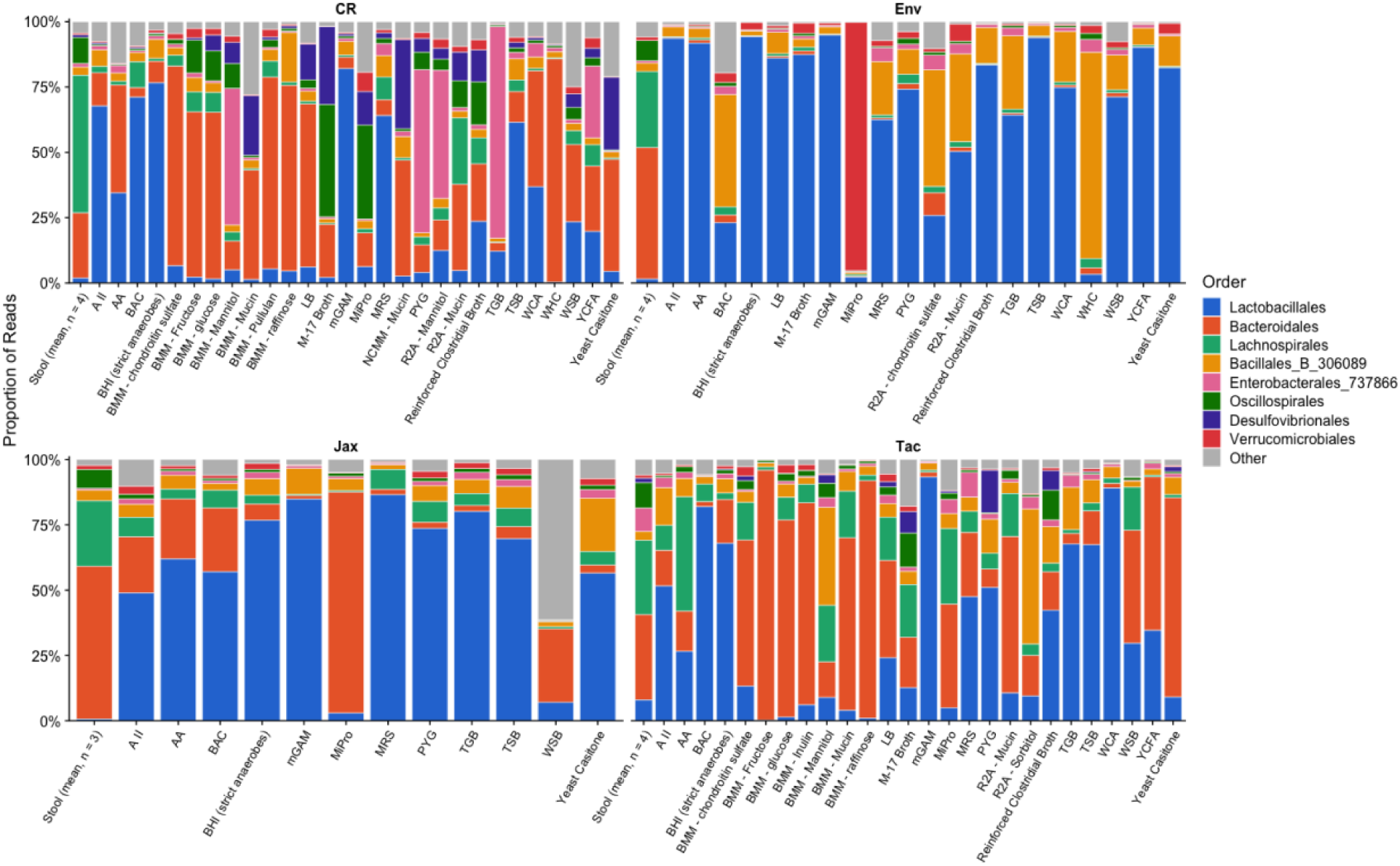
Taxonomic composition shifts from stool to culture across media conditions. Barplot of read relative abundance of Bacteria at taxonomic Order level for each media condition supporting growth. Stool replicates are averaged. Stool communities are dominated by *Bacteroidales*, *Lachnospirales*, and *Oscillospirales*. In culture, *Lactobacillales* dominates many conditions, while BAC cultures show very low diversity, illustrating the taxonomic shift from stool to culture and the gradient of niche selectivity across media. Taxonomy was assigned using GTDB and collapsed to 160 organisms.

**Supplementary Figure 5.**
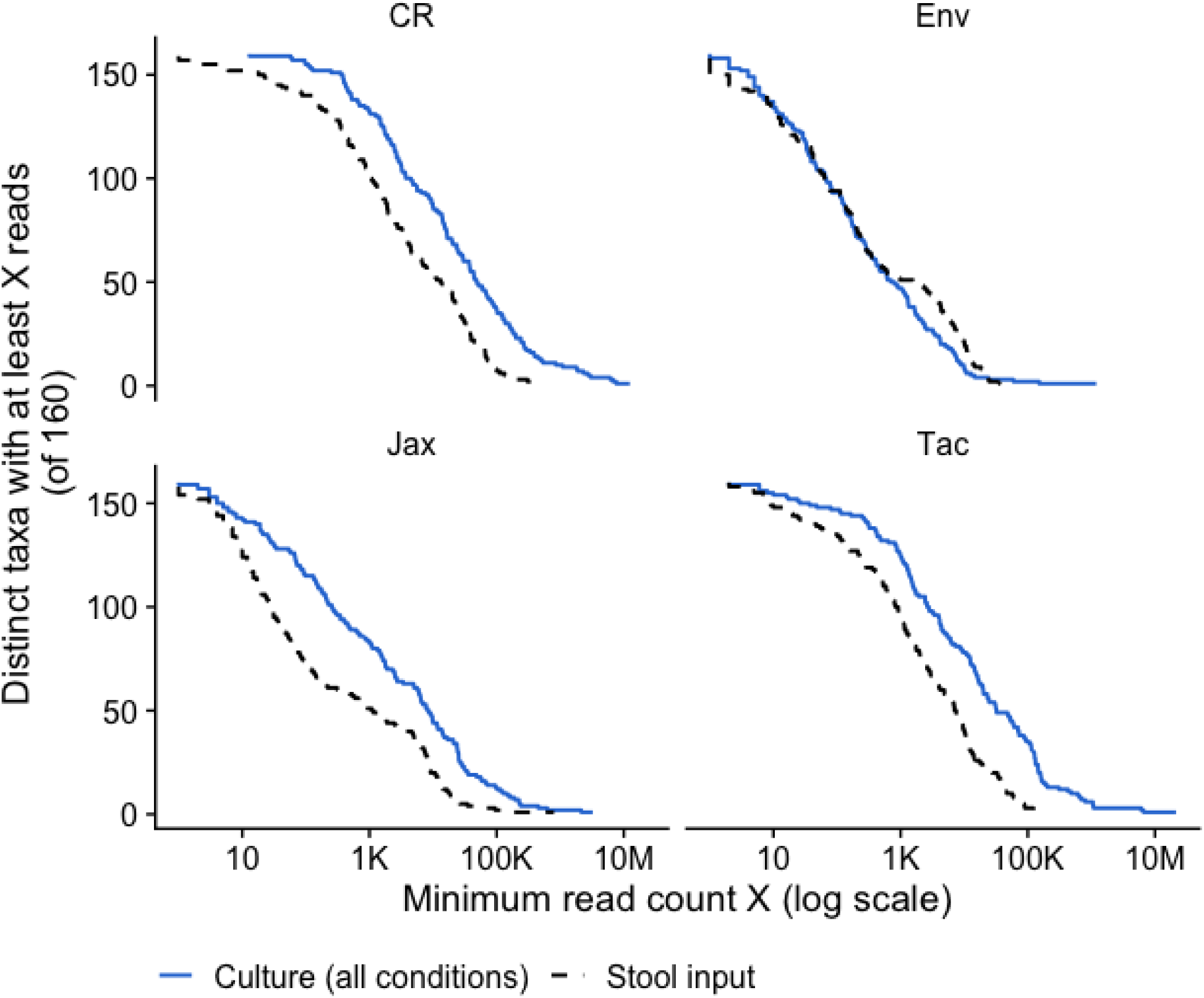
The 60-medium panel recovers nearly all stool-detected taxa. Line plot showing the number of organisms (of 160 collapsed organisms) detected at or above increasing minimum read count thresholds, comparing pooled culture recovery to stool input for each vendor. At a threshold of 10 reads, nearly 100% of stool taxa are recovered across all vendors, with recovery declining at higher stringency. This demonstrates that the diversity of media conditions in the panel is sufficient to capture the vast majority of cultivable stool diversity. Recovery was assessed by pooling detections across all 60 media conditions per vendor.

**Supplementary Figure 6.**
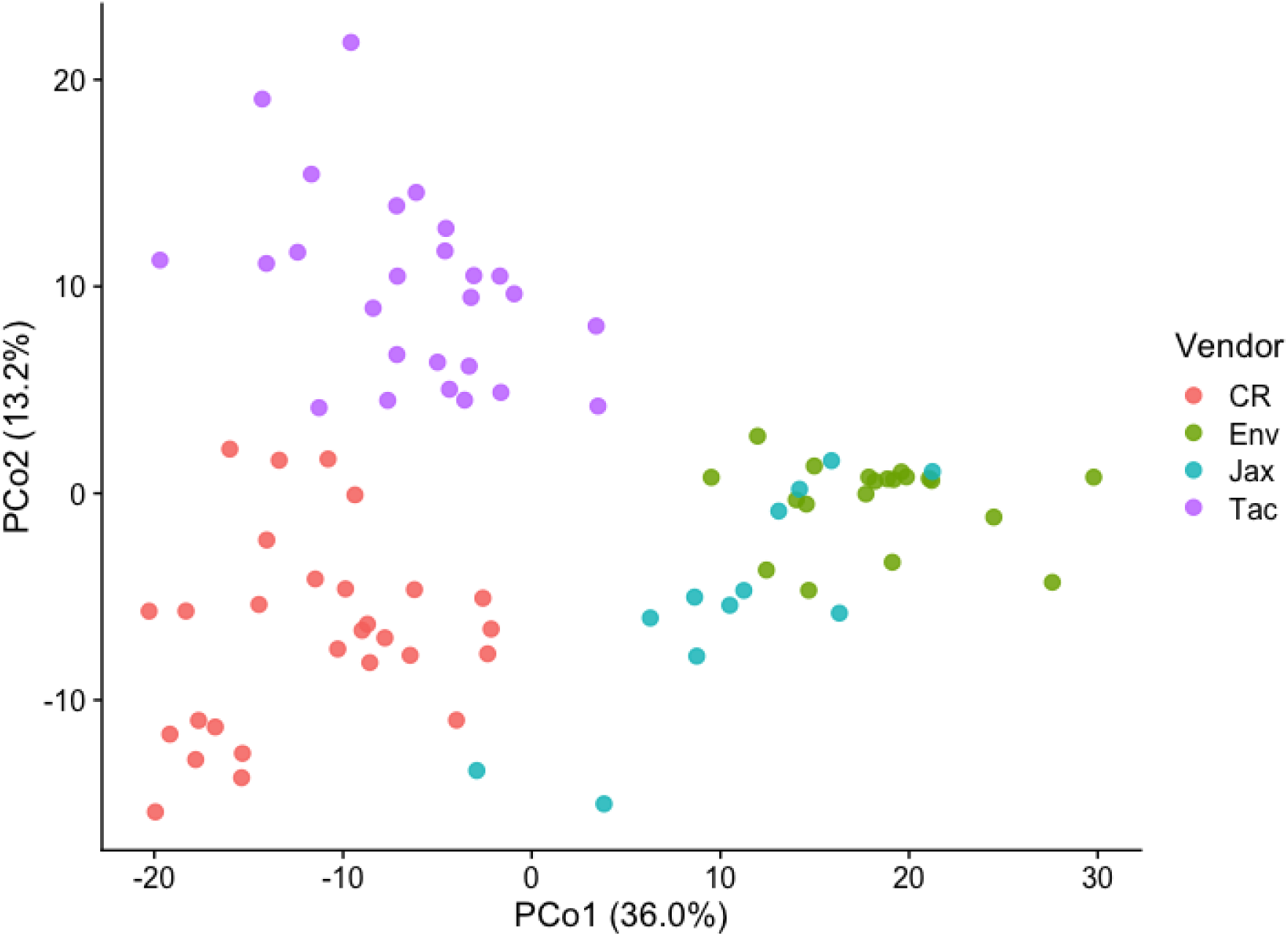
Cultured communities cluster by vendor of origin. PCoA of Aitchison distances (Euclidean on CLR-transformed counts) across 86 QC-passing culture libraries, colored by mouse vendor. PCo1 captures 36.0% and PCo2 13.2% of variation. CR and Tac separate clearly, while Env and Jax partially overlap. This indicates that starting community composition is the dominant driver of cultured community structure, even across diverse media conditions. Libraries were filtered to OD_600_ > 0.1 above blank and >= 100,000 reads. The distance matrix was computed from 160 collapsed organisms.

**Supplementary Figure 7.**
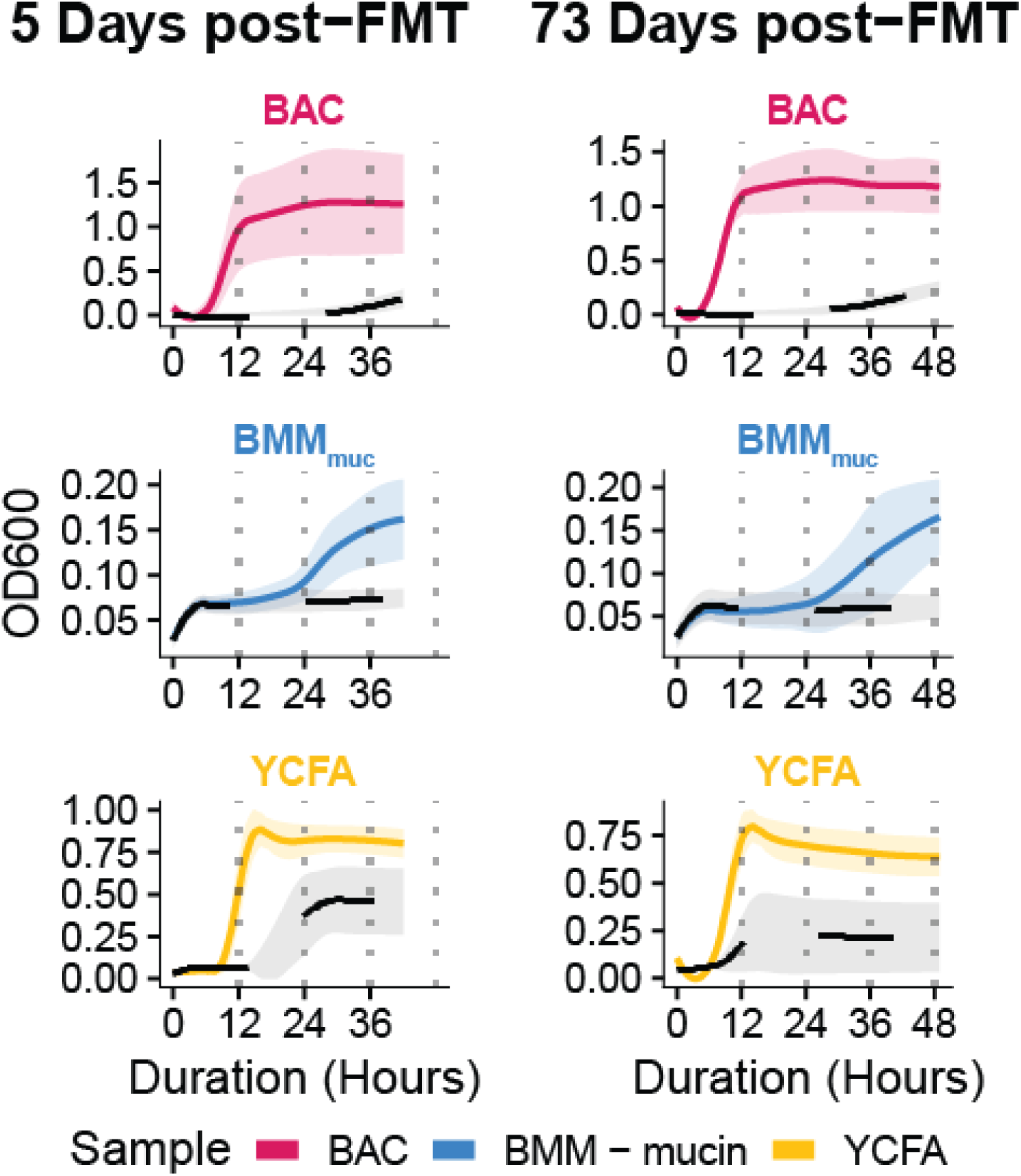
Transplanted NACs engraft in mice and persist long term. Growth curves of stool samples from mice gavaged with a specified NAC or PBS, collected at day 5 or day 73 post-gavage. Samples were cultured in the same medium used to propagate the corresponding NAC. Data represent the mean of nine replicates; shaded areas indicate standard error of the mean.

